# Investigating the role of N-terminal domain in phosphodiesterase 4B-inhibition by molecular dynamics simulation

**DOI:** 10.1101/2020.05.12.090993

**Authors:** Vidushi Sharma, Sharad Wakode

**Affiliations:** Department of Pharmaceutical Chemistry, Delhi Institute of Pharmaceutical Sciences & Research, Mehrauli-Badarpur Road, Pushp vihar Sector 3 New Delhi 110017, India

**Keywords:** Phosphodiesterase 4B, UCR2, Molecular dynamics simulation, Principal component analysis, Distance cross correlation matrix

## Abstract

Phosphodiesterase 4B (PDE4B) is a potential therapeutic target for the inflammatory respiratory diseases such as congestive obstructive pulmonary disease (COPD) and asthma. The sequence identity of ∼88% with its isoform PDE4D is the key barrier in developing selective PDE4B inhibitors which may help to overcome associated side effects. Despite high sequence identity, both isoforms differ in few residues present in N-terminal (UCR2) and C-terminal (CR3) involved in catalytic site formation. Previously, we designed and tested specific PDE4B inhibitors considering N-terminal residues as a part of the catalytic cavity. In continuation, current work thoroughly presents an MD simulation-based analysis of N-terminal residues and their role in ligand binding. The various parameters viz. root mean square deviation (RMSD), radius of gyration (Rg), root mean square fluctuation (RMSF), principal component analysis (PCA), *dynamical cross-correlation matrix* (*DCCM*) analysis, secondary structure analysis, and residue interaction mapping were investigated to establish rational. Results showed that UCR2 reduced RMSF values for the metal binding pocket (31.5±11 to 13.12±6 Å^2^) and the substrate-binding pocket (38.8±32 to 17.3±11 Å^2^). UCR2 enhanced anti-correlated motion at the active site region that led to the improved ligand-binding affinity of PDE4B from −24.57±3 to −35.54±2 kcal/mol. Further, the atomic-level analysis indicated that T-pi and π-π interactions between inhibitors and residues are vital forces that regulate inhibitor association to PDE4B with high affinity. In conclusion, UCR2, the N-terminal domain, embraces the dynamics of PDE4B active site and stabilizes PDE4B inhibitor interactions. Therefore the N-terminal domain needs to be included by designing next-generation, selective PDE4B-inhibitors as potential anti-inflammatory drugs.

## 1. Introduction

Respiratory inflammatory diseases such as congestive obstructive pulmonary disease (COPD) and asthma affect millions worldwide(Zhang, Ibrahim, Gillette & Bollag 2005). Phosphodiesterase 4B (PDE4B) is a potential therapeutic target in inflammatory diseases. It hydrolyzes cyclic adenosine monophosphate (cAMP), an important secondary messenger, to 5’-adenosine monophosphates (AMP). AMP is a potent pro-inflammatory mediator in humans, and thus PDE4B regulates many biological processes inside the cell by regulating the production of AMP (Houslay & Adams 2003). Therefore, PDE4B inhibitors that prevent cAMP hydrolysis are much needed. The PDE4 enzyme family consists of four members (named PDE4A, PDE4B, PDE4C, and PDE4D), and the mechanism of cAMP catabolism is conserved among all the four PDE4 enzymes. These PDE4 enzymes are highly expressed in neutrophils, monocytes, central nervous system, and smooth muscles of the lung. Available non-selective PDE4 inhibitors like rolipram cause severe side effects such as nausea and emesis. As an improvement, the second-generation PDE4 inhibitors like roflumilast show fewer side effects; however, they are associated with a narrow therapeutic window. Knockout studies in mice suggest that available PDE4B inhibitors co-inhibit PDE4D, a close homologous enzyme, thus causing severe side effects (Wang et al. 1997; Calverley et al. 2009). Therefore, selective inhibition of PDE4B is much needed to develop safe anti-inflammatory drugs. Nonetheless, the high sequence (and thus structural) similarity between the two enzymes, PDE4B and PDE4D, sets hurdles to design and develop selective PDE4B inhibitors.

The structural comparison of the two enzymes, PDE4B and PDE4D, reveals that the active site cavity is composed of residues from the catalytic domain and a regulatory domain. These regulatory domains include the N-terminal, upstream conserved region 2 (UCR2) domain, and the C-terminal regulatory domain (Rocque et al. 1997). Either of these two regulatory domains can cover the catalytic cavity, and thus, provide selectivity for PDE4B inhibition. Contrary to the role of UCR2 in PDE4B inhibition, the UCR2 domain does not affect the substrate affinity (Km) of PDE4B, as shown by similar Km values for the ‘long-’, and the ‘catalytic-’ versions of PDE4B (Saldou et al. 1998). In this work, we have focused on studying the structural effect of the regulatory N-terminal domain, UCR2, on PDE4B inhibition. While a catalytic domain is conserved, with an RMSD value of 0.352 (Cα atoms; PDE4B:PDE4D::3G45:3G4G), between the two enzymes, the UCR2 domain shows following distinct residues that cover the active site region: PDE4B:PDE4D::Tyr274:Phe196 (Rocque et al. 1997). Burgin *et al*. have established that Tyr274 indeed provides selectivity for inhibition of PDE4B. Although, other groups and we have reported selective PDE4B inhibitors, and evaluated the energetics of PDE4B inhibition (Goto et al. 2014; Li et al. 2016; Sharma & Wakode 2017; Xing, Akowuah, Gautam & Gaurav 2017), yet, the structural dynamics of PDE4B inhibition was merely understood. Understanding the dynamics of UCR2-domain guided PDE4B inhibition will guide us to design and optimize available PDE4B inhibitors. Literature supports that similar MD simulation studies can successfully be implemented to understand the mechanisms of molecular recognition and conformational changes in therapeutic targets (Padhi, Kumar, Vasaikar, Jayaram & Gomes 2012; Sharma & Wakode 2016; Moreau et al. 2017; Santos et al. 2017; Douglas et al. 2018; Elfiky 2020; Islam et al. 2020; Kumar et al. 2020). In this work, we performed a comparative all-atom molecular dynamics simulation study of PDE4B-inhibitor complex and explored the effect of UCR2-domain residues in PDE4B inhibition. In 100ns long MD simulations, we performed principal component analysis (PCA), dynamical cross-correlation matrix (DCCM) analysis, secondary structure analysis, and residue network mapping to shed light on the fact that the N-terminal UCR2 domain improves non-polar interactions at PDE4B-inhibitor interface. Therefore, these UCR2 domain residues must be targeted to optimize PDE4B inhibitors (Tautermann, Seeliger & Kriegl 2015; Xiao, Yang, Xu & Vongsangnak 2015).

## 2. Experimental

### 2.1 Data selection

The highest resolution PDE4B crystal structure [PDB ID-2QYL inhibitor-NPV (4-[8-(3-nitrophenyl)-1,7-naphthyridin**-**6-yl] benzoic acid) resolution: 1.95 Å] was retrieved from the protein data(Berman et al. 2000). To study the role of the N-terminal UCR2 domain, we retrieved another PDE4B structure (PDB ID: 3G45, inhibitor: NPV, 8-(3-nitrophenyl)-6-(pyridin-4-ylmethyl)quinoline and resolution: 2.63 Å). Here onward, these complex structures will be denoted as “**U-Cat”** (structure with **<u>U</u>**CR2+**<u>Cat</u>**alytic domain; PDB ID: 3G45) and “**Cat”** (structure with only **<u>cat</u>**alytic domain; PDB ID: 2QYL). To understand the role of UCR-2 residues in PDE4B-inhibitor interactions, we used a common inhibitor, NPV, in both the structures and replaced the coordinates of 988, a co-crystallized ligand, with NPV in the 3G45 structure.

### 2.2 Protein Preparation

Previously retrieved structures may have missing bond order, connectivity, steric clashes or bad contacts with the neighboring residues. Therefore, both the structures were prepared using Protein Preparation Wizard (Impact 6.3, Schrodinger 2014-2, Maestro 9.8) (Sastry, Adzhigirey, Day, Annabhimoju & Sherman 2013) as previously described (Sharma & Wakode 2016). In brief, both the structures were corrected for atoms and bonds, and energy minimized to potentially relax the structures to get rid of any steric clashes in the structures.

### 2.3 Molecular Dynamic Simulations

FF14SB force field parameters were set for the protein using the AMBER14 (Assisted Model Building with Energy Refinement) LeaP module (Hornak et al. 2006). AM1-BCC method was used to assign partial atomic charges for bound inhibitor (NPV), and the general amber force field (GAFF) was used to create its topology (Wang, Wolf, Caldwell, Kollman & Case 2004). Mg^2+^ and Zn^2+^ ions were treated according to the “non-bonded” model method (Stote & Karplus 1995). The prepared systems were solvated with the TIP3P water model by creating a cubic water box, where the distance of the box was set to 10Å from the periphery of protein (Kiss & Baranyai 2011). Molecular systems were neutralized through the AMBER LeaP module by adding the necessary amount of counter ions (Na^+^) to construct the system in an electrostatically preferred position.

The whole assembly was then saved as the prepared topology and coordinate files to use as input for the PMEMD module of the AMBER (Case et al. 2005). At the first step, the prepared systems were energy minimized in a two-step process: initial 1000 steps of steepest descent followed by 500 steps conjugate gradient. In the first part of minimization, the complex was kept fixed to allow water and ion molecules to move, followed by the minimization of the whole system (water, ions and complex) in the second part. The minimized systems were gradually heated from 0 to 298 K using a NVT ensemble for 100ps where the protein-ligand complex was restrained with a large force constant of 5 kcal/mol/Å^2^. Following heating, the systems were equilibrated under constant pressure at 298 K, and the restrain was gradually removed at NPT ensemble as follows: 5 kcal/mol/Å^2^ (40 ps), 2 kcal/mol/Å^2^ (20 ps), 1 kcal/mol/Å^2^ (20 ps) and 0.5 kcal/mol/Å^2^ (10 ps). Final simulations, the production run, were performed for 100 ns on NPT ensemble at 298 K temperature and 1 atm pressure. The step size of 2 fs was kept for whole simulation study. Langev in thermostat and barostat were used for temperature and pressure coupling. SHAKE algorithm was applied to constrain all bonds containing hydrogen atoms (Gunsteren & Berendsen 2006). The non-bonded cutoff was kept on 10Å, and long-range electrostatic interactions were treated by Particle Mesh Ewald method (PME) with a fast Fourier transform grid spacing of approximately 0.1nm (Darden, York & Pedersen 1993). Trajectory snapshots were taken at each 100ps, which were used for final analysis. The minimization and equilibration were performed by the PMEMD module of AMBER14. The production simulations were performed using the PMEMD program of AMBER running on NVIDIA Tesla C2050 GPU work station (Gotz et al. 2012). The production run was considered for the analysis which was carried out using the cpptraj module of the AMBER14 and VMD (Humphrey, Dalke & Schulten 1996)

### 2.4 RMSD

The stability of the two systems during the simulations was studied by calculating RMSD of the backbone atoms of different frames to the initial conformation and therefore, RMSD is the measure of the average distance between the atoms (usually the backbone atoms) of superimposed protein structures. RMSD of Cα atoms was calculated using the cpptraj analysis tool in the AMBER 14 program.

### 2.5 Rg

The radius of gyration is a measure of the distance between the center of mass and both termini of the protein and therefore radius of gyration determines the compactness of protein structure (Lobanov, Bogatyreva & Galzitskaia 2008). The average Rg was computed by taking the average of Cα atoms over at least 5000 frames of the trajectories.

### 2.6 RMSF

RMSFs were calculated with the backbone atoms of amino acid residues for both the trajectories using the cpptraj module of AMBER 14. The starting conformations of each complex structure were used to align the coordinates of the two trajectories.

### 2.7 DCCM

The cross-correlation was calculated as block average over time from 50 ns to 100 ns from the MD trajectories of the two systems, U-Cat and Cat. We use the first coordinate set of the analysis portion of the simulation, *i.e.*, starting structure as a reference set. Each subsequent coordinate set was translated and then rotated to obtain the minimum RMS deviation of the Cα atoms from the reference coordinate set (McCammon & Harvey 1988).

### 2.8 PCA

PCA was performed using the cpptraj module of AMBER 14 to understand the collective atomic motion of U-Cat and Cat versions of PDE4B. Atomic coordinates extracted from the last 50 ns trajectories to study the covariance matrix of the two systems. The eigenvalues and the projections along the first three PCs were calculated as previously described (Amir et al. 2019; Fatima et al. 2019).

### 2.9 Protein secondary structure analysis

In the dynamic analysis, it is imperative to identify critical changes in the secondary structure element during the simulation. It imparts deep insight into the stability of the secondary structure. Each secondary structure type is shown by a different color, and a change in secondary structure type is easily differentiated. The secondary structure information for each frame was calculated, and a two-dimensional plot was generated for the last 50 ns trajectory, 500 frames.

### 2.10 Residue network mapping

The study of residue interacting patterns is imperative to analyze protein’s structural rigidity, secondary structure maintenance, and functionality. It includes crucial interactions, hydrogen bond occupancies that are present during the simulation. These interactions are identified using VMD. The hydrogen bond interaction was calculated between the polar hydrogen atom and a nearby (< 3.4 Å) acceptor atom. Similarly, the π -π interactions were calculated in the crystal structure and MD-simulated structure of the last 100th ns. The π -π interactions results with the attraction of one π electron cloud system with another nearby π electron cloud system. These interactions are crucial in bioprocesses. In this study, we measure the inter-residue distance to each hydrophobic ring centroid to identify the pair of π -π interacting residues. The residue pairs within a ring centroid distance of 3.0–7.5 Å (min-max) were considered. Further, we calculated the dihedral angle (θ) between the two planes that have two types of structural geometries. If the dihedral angle between the planes of two rings was found to be 30° > θ > 150°, the rings are in face-to-face interaction and if the angle is between 30° < θ < 150°, the rings are in T-shaped (edge to face or perpendicular) interactions (Anjana et al. 2012).

## 3. Result and discussion

The objective of our study was to understand the role of N-terminal UCR2-domain residues in the stability of PDE4B-inhibitor complexes. To this end, we performed all-atom molecular dynamics simulations of PDE4B-NPN complex in the presence (**U-Cat**) and absence of the UCR2-domain residues (**Cat**). First, we confirmed the stability of two systems by RMSD analysis and total energy calculations. Next, the stable MD trajectories were processed to understand the dynamics of PDE4B-NPV interactions through RMSF analysis, DCCM analysis, and PCA. Lastly, we analyzed residue network mapping.

### 3.1 Systems stability

The reliability of our MD simulations was assessed in terms of RMSD values, total energies, and radius of gyration Rg of the two simulations as follows:

#### 3.1.1 RMSD

RMSD analysis accounts for the average distance between the selected atoms of superimposed biomolecules, which indicates the closeness between the three-dimensional structures. In this study, we calculated the RMSD values for the backbone atoms for the proteins and all non-hydrogen atoms for the small molecule, NPV. Within the initial 5-ns of each simulation, both the systems became stable and remained stable throughout the 100ns long MD simulations **(Fig. 1)**. During the last 50 ns long simulations, the RMSD values of catalytic domain residues (Ser324-Pro665) were as follows: 1.28 ± 0.12 Å (**Cat**) and 1.11 ± 0.09 Å (**U-Cat**). The lower magnitude of these values suggests the stability of both the systems. Further, the difference between these values for both the systems (**U-Cat** versus **Cat** complexes) is insignificant (t-test). This finding suggests that the additional N-terminal, UCR2-domain does not affect the major dynamics of the catalytic domain of the protein.

**Figure 1:**
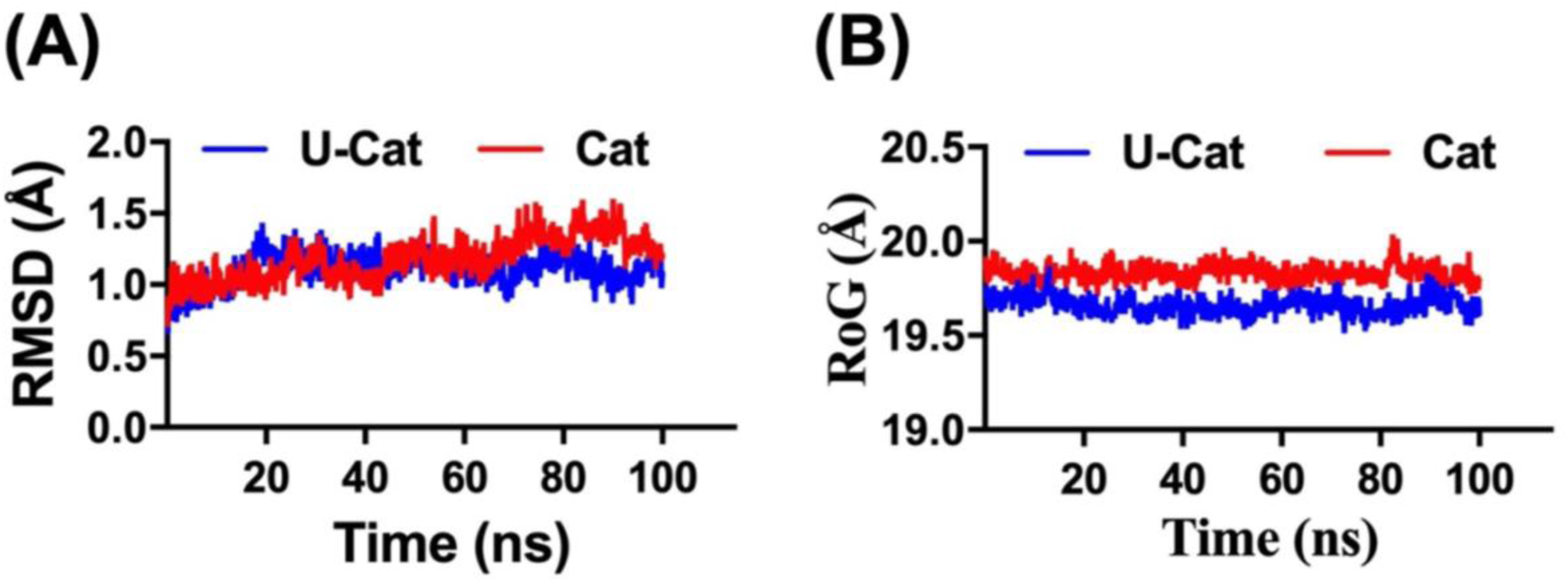
RMSD and Rg plots of two complexes.

Similar to RMSD values for the catalytic domain, the RMSD values for the bound inhibitor, NPV, are also low (<1.5 Å), suggesting the stable dynamics of the two systems. The RMSD values for the two systems during the last 50 ns duration are as follows: 1.16±0.07 Å (**Cat**) and 0.86±0.26 Å (**U-Cat**). The lower RMSD values for the **U-Cat** system suggest that additional UCR2-domain stabilizes the bound inhibitor.

#### 3.1.2 Rg

Rg measures the mass-weighted RMS distance of a group of atoms from their common center of mass, which indicates the global dimension of the protein. For both the systems (**Cat** and **U-Cat**), we calculated the Rg for the catalytic domain residues (Ser324-Pro658) and observed a stable trend in Rg plot. The Rg values for the last 50 ns simulations for the two systems were as follows: 19.66±0.05 (**U-Cat** system) and 19.78 ± 0.05 (**Cat** system) (**Fig. 1B**).

### 3.2 Structural dynamics of PDE4B-NPN complex in the presence (U-Cat) and absence (Cat) of UCR2-domain

Once we confirmed the stability of the two systems, next we sought to study the conformational changes in the two complexes (**Cat** and **U-Cat**) and performed the following analyses:

#### 3.2.1 RMSF

To understand and compare the dynamics of the two complexes, we calculated the RMSF values for each complex. Using RMSF analysis, we studied the fluctuation at the individual residue level in the two systems. To compare the two systems, we rejected the additional N-terminal residues in **the U-Cat** structure and focussed on the catalytic domain residues in the two complex structures. The comparative RMSF plot of the two systems is shown in **Fig. 2**. A similar trend in RMSF values suggested that the UCR2-domain had no drastic effect on the conformational dynamics of the two systems.

**Figure 2:**
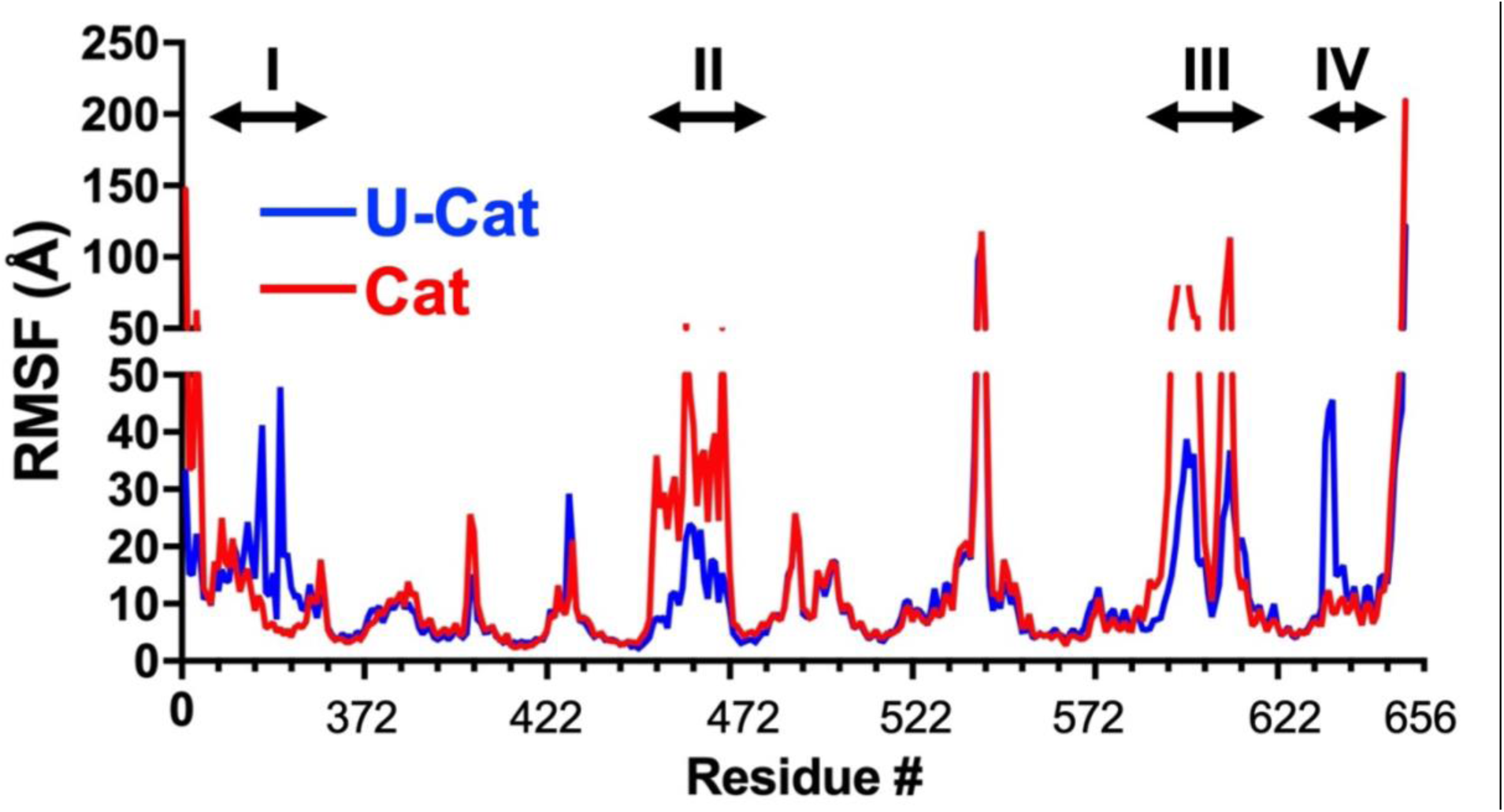
RMSF plot of the two complexes.

Intriguingly, we observed the distinct behaviors of the two complexes, **U-Cat** and **Cat**, in following four regions: Asp336-Asn354 (region I; 19 residues), Pro451-Glu472 (region II; 22 residues), Phe587-Ser614 (region III; 28 residues) and Gln635-Asp643 (region IV; 13 residues). Interestingly, region II residues belong to the metal-binding pocket such as His450, Asn283, Glu476. The catalytic residue, Gln615, and the P clamp pocket (Met583, Met603, and Val611) belong to region III. Region I and IV are distant from the active site region and do not include any catalytic or functional residue. Among these regions, the presence of additional UCR2-domain residues stabilized the region II and III in **the U-Cat** complex, which was reflected in lower RMSF values as compared to **Cat** complex. In contrast, the **region I** and **IV** showed higher RMSF values for the **U-Cat** complex.

Based on the locations of these four regions to **UCR2-domain**, we hypothesized that UCR2-domain might affect the associated motion of the catalytic domain of PDE4B and sought to perform the dynamical cross-correlation matrix analysis.

#### 3.2.2 DCCM

To understand the variation in the RMSF values of the two complexes and to check if motions in different regions (I-IV) are related or not, we performed an inter-residue DCCM analysis for Cα atoms to find out the extent of correlation of atomic movements (**Fig. 3**). To compare the two complex structures, **U-Cat** and **Cat**, we only considered the catalytic (**Cat**) residues in the two complexes to perform the DCCM analysis. Similar diagonal trends in the two figures suggested that the UCR2-domain had a severe effect on the secondary structure of the two complexes suggesting that the two systems, **U-Cat** and **Cat**, may have similar global dynamics. Further, the darker shades of colors in the **U-Cat** complex suggested that the presence of additional UCR2-domain caused a greater extent of correlative (green) and anticorrelated (purple) motions in PDE4B-NPV complex. In particular, we observed intense purple shades in regions II and III of the **U-Cat** complex suggesting more substantial anticorrelated motions in these regions. Intriguingly, these anti-correlative motions in regions II and III were correlated with low RMSF values in these regions (**Fig. 2 and 3**). On the contrary, the region I and IV showed enhanced correlative motion in the presence of UCR2-domain in the **U-Cat** complex. In conclusion, the darker shades plausibly suggest the enhanced associated motion U-Cat complex. Furthermore, low RMSF values and enhanced anticorrelated motion in regions II and III suggest that the presence of the UCR2 domain stabilizes the PDE4B-inhibitor complex through enhanced anticorrelated motion (**Fig. 2 and 3**). Interestingly, these results were in line with the energetics of the two systems because, in our MM-GBSA analysis, we observed the U-Cat complex’s enhanced stability compared to the Cat-complex (**Supplementary data**).

**Figure 3:**
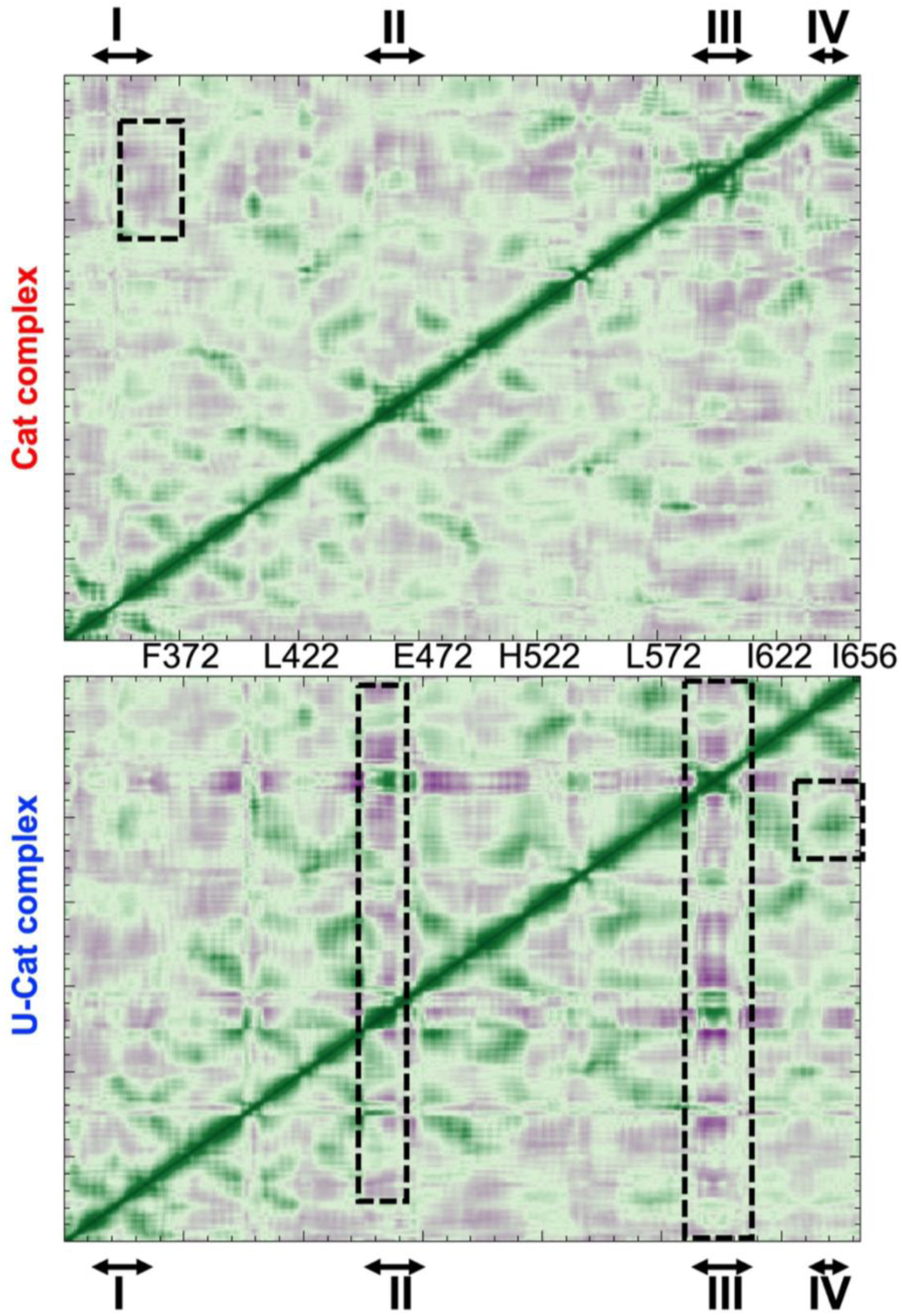
DCCM maps of the two complexes.

#### 3.2.3 PCA

Through the DCCM study, we confirmed that the presence of UCR2-domain affected the correlative (and anti-correlative) motions in PDE4B-NPV complex. To understand the directionality of these motions, we proceeded further and performed a PCA of the two simulations (**U-Cat** and **Cat**). Here, we designated the whole average of receptor motion based on Cα atoms. This approach supported to figure out the overall combined motion of the Cα atoms in the protein denoted by the eigenvectors of the covariance matrix, which is asserted by its coincident eigenvalues (Amadei, Linssen & Berendsen 1993). The occurrence of the eigenvectors associated with large eigenvalues can generally represent the over-all concerted motion of the protein correlated to the protein function. The increasing sum of eigenvalues as a function of the number of eigenvalues resulting from the MD simulation frames during 0.1μs is shown in **(Fig. 4A)**. To compare the two complexes, we ruled out additional residues of UCR2-domain in **U-Cat**. We observed that the first two components of the analysis, PC1, and PC2 were significantly higher for the **U-Cat** complex than the **Cat** complex, suggesting that the additional UCR2-domain enhanced the directional motion at the catalytic domain of PDE4B. **Fig. 4B** shows the domains, directions, and degrees of motions corresponding to the first three PCs of each complex. In all three projections, **the U-Cat** complex occupied a broad range of phase spaces as compared to **the Cat** complex. An increase in the collective motions of **U-Cat** may plausible enhanced interaction and thus the stability of the complex as compared to **Cat** complex. We next studied directionality and magnitudes of the three PCs using porcupine plots. The changes in direction and magnitude of **U-Cat** suggested that the additional UCR2-domain posed a significant conformational impact on the overall dynamics of PDE4B. We found that PC1 and PC2 motions were confined to regions Phe587-Ser610 and Asp447-Asp472 in the **Cat** complex. On the contrary, the motion in the active site region decreases in the **U-Cat** complex suggesting that the additional UCR2-domain indeed stabilizes the catalytic cavity region in PDE4B (**Figure 4**).

**Figure 4.**
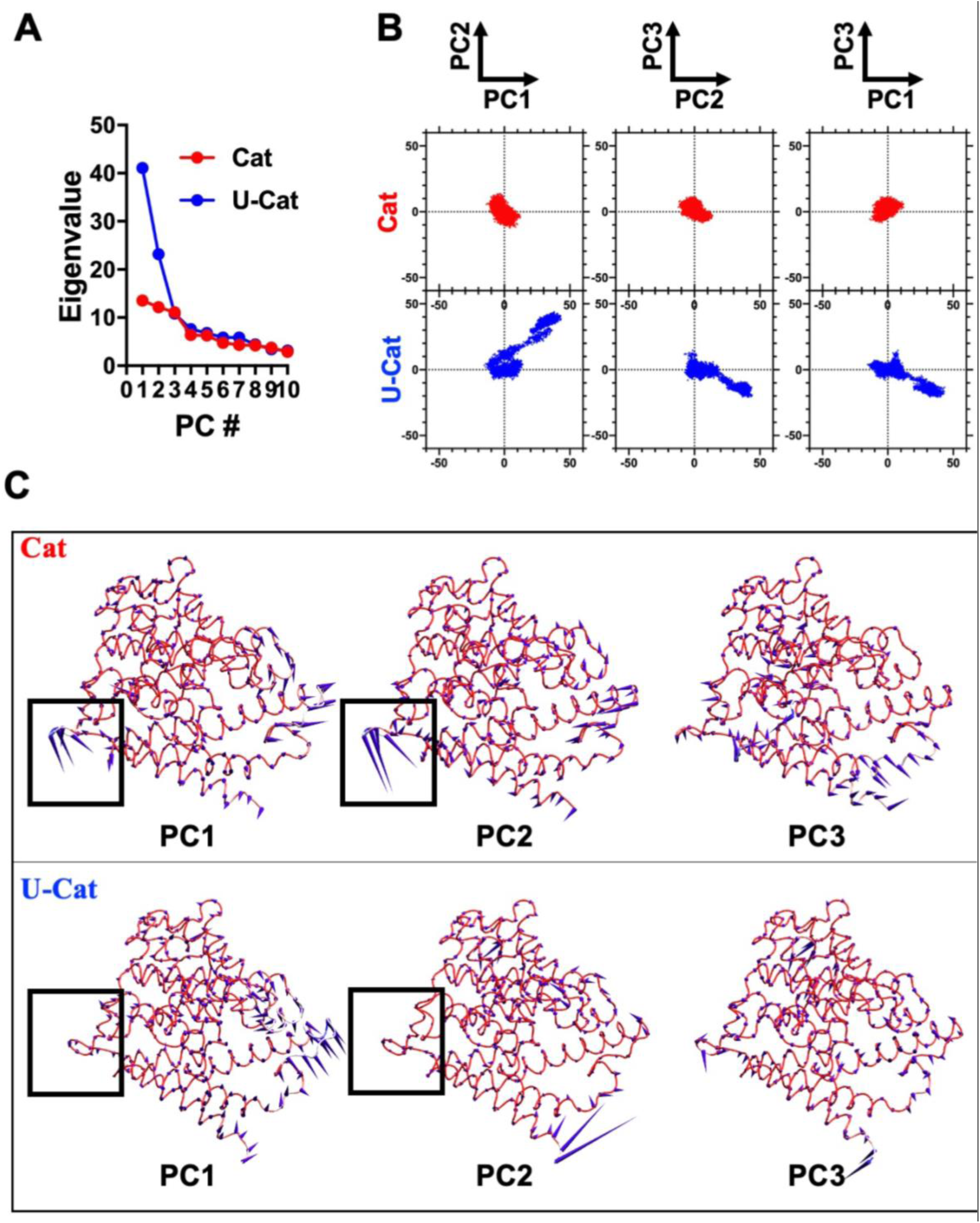
PCA of the two complexes. Top(left): Plot of first-ten principal components (PCs). Top (right): Plot of first three-PCs. Bottom: Porcupine plots of first 3-PCs of the two complexes. Please note that in the U-Cat complex, the motion at the active site region decreases as highlighted by the squares.

To conclude, our RMSF, DCCM, and PCA results suggested that the presence of UCR2-domain affected the dynamics of the catalytic domain of PDE4B. Interestingly, region II and III that showed enhanced anticorrelated motions and reduced RMSF at the active site region. On the contrary, the region I and IV that showed enhanced correlative motion and larger RMSF values, but they were distant from the active site region. Therefore, the additional domain UCR2-domain enhances the anti-correlative motion at the active site cavity and thus stabilized the protein-inhibitor interactions.

#### 3.2.4 Secondary structure analysis

Further, to visualize critical changes in the protein conformation during the timeline analysis, we performed a secondary structure analysis of the two simulations. We generated a two-dimensional plot secondary structure information for the 100ns trajectory (**Fig. 5**). We observed that region Ser423-Asp433, which was a turn in the **U-Cat** complex, changed to a 3-turn helix in the **Cat** complex (brown to green). Further, we observed a local structural transition at the region Glu343–Phe353 that was a turn in the **Cat** complex but transformed into a coil in the presence of the UCR2-domain in the **U-Cat** complex, shown as green to red transition in **Fig. 5**. We observed a similar transition at the region Thr535-Thr550 that turned from a coil in the **Cat** complex to an anti-parallel beta-sheet in **the U-Cat** complex as seen a transition from red to blue color in the **Fig. 5**. Intriguingly, these differences in the secondary structure were in agreement with the RMSF and DCCM analyses and thus suggested the flexible residues.

**Figure 5.**
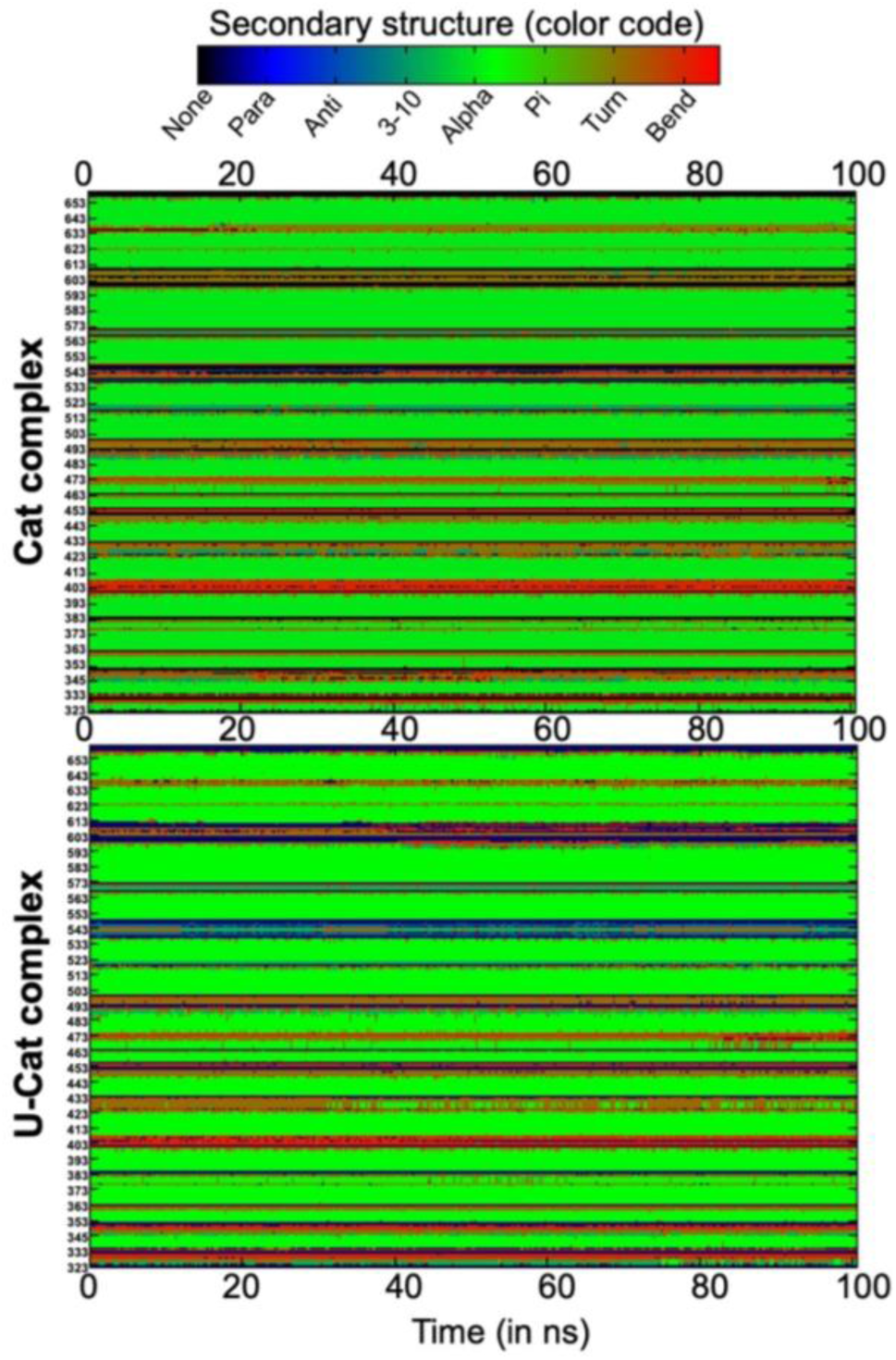
Secondary structure plot of the two complexes during 100 ng long simulations.

### 3.3 The dynamics of the active site region

To further understand the effect of additional UCR2-domain on the catalytic domain of PDE4B, we studied the dynamics of the active site region in detail. At first, we assessed the stability of an H-bond with Gln615, an interaction reported in the crystal structure. We observed a stable H-bond with Gln615 in both the simulations, **U-Cat**, and **Cat** complexes. Further, we observed an enhanced occupancy of 65% of Gln615-H-bond in the **U-Cat** complex compared to 43% observed in **the Cat** complex, suggesting that the additional UCR2-domain indeed stabilizes the PDE4B-inhibitor interactions.

Furthermore, we observed an additional H-bond with Tyr274 (occupancy 25%) in **the U-Cat** complex, which was otherwise absent in the simulation of the Cat complex. Next, we studied the dynamics of the other plausible H-bonding residues in the active site cavity (Sharma & Wakode 2017). These residues include Tyr274, Tyr405, His406, His410, Tyr415, Asp447, Asp564, Asn567, and Gln615. We observed that these plausible H-bonding residues were more stable in the **U-Cat** complex as compared to **the Cat** complex as interpreted by lower RMSD values.

In addition to H-bonding residues, π-π interactions with Phe618, Phe586, Tyr405, and Tyr274 were stable and consistent with the interactions reported in the crystal structures (**Fig. 6**). To this end, the MM-GBSA analysis also confirmed the enhanced stability of the **U-Cat** complex as compared to the Cat-complex (**Supplementary data**).

**Figure 6.**
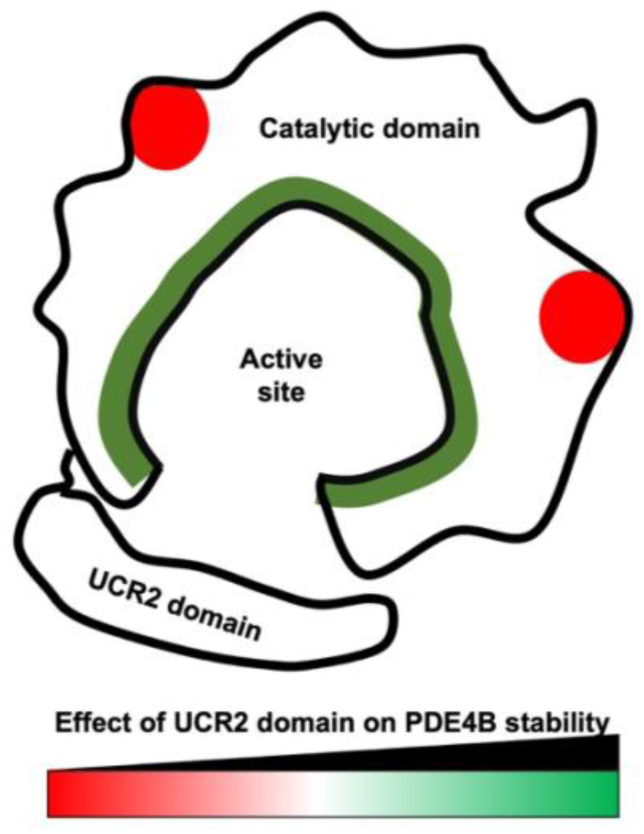
An illustration to show the effect of additional UCR2 domain residues on the stability of the catalytic domain.

Altogether, our simulations and post-trajectory analyses confirm that the additional UCR2-domain stabilizes the PDE4B-inhibitor complexes and therefore, must be considered while evaluating the binding stability of newly designed PDE4B inhibitors.

## 4. Conclusion

We performed 100 ns long molecular dynamics simulations of PDE4B variants, with and without the UCR2-domain. Both complexes with a common inhibitor, NPV, and probed the effect of additional UCR2-domain on the stability of PDE4B-NPV interactions. RMSF, DCCM, PCA, H-bond network analyses confirm that the UCR2-domain enhances the correlation and anticorrelation motion in the complex structure. This study provides substantial evidence to include the regulatory UCR2-domain in further drug designing to evaluate the stability of newly designed PDE4B inhibitors.

## List of abbreviations

PDE4B: Phosphodiesterase 4B
cAMP: Cyclic adenosine monophosphate
COPD: Chronic obstructive pulmonary disease
PDB: Protein data bank
MD: Molecular dynamics
RMSD: Root mean square deviation
RMSF: Root mean square fluctuation
DCCM: dynamical cross-correlation matrix DCCM
PCA: Principal component analysis
U-cat: PDE 4B structure containing UCR2, and catalytic domain
Cat: PDE 4B structure containing the catalytic domain

