## Supplemental data for "Investigating the role of N-terminal domain in phosphodiesterase 4B-inhibition by molecular dynamics simulation"

### MM-GBSA calculations

#### Method

The binding affinity of PDE4B-inhibitor complexes are studied using binding free energy calculations which were estimated using the MM-GBSA (Molecular Mechanics Generalized Born Surface Area) method (Genheden & Ryde 2015). Additionally, different components of the interaction energy that contribute to the binding were estimated. The entropy was not calculated, as we were interested in the comparative analysis of interaction parameters important to the inhibitor binding. Binding free energies ($\Delta G_{Bind}$) of Ligands at catalytic cavity of protein was calculated as follows:

$$\Delta \boldsymbol{G}_{\boldsymbol{Bind}} = \boldsymbol{G}_{\boldsymbol{Complex}} - \boldsymbol{G}_{\boldsymbol{Protein}} - \boldsymbol{G}_{\boldsymbol{Ligand}} \qquad \textbf{(1)}$$

Where, $G_{Complex}$ $G_{Protein}$ and $G_{Ligand}$ is the free energy of complex, protein (PDE4B), and ligand, respectively. Free energy (ΔG) of each state was calculated as follows:

$$\boldsymbol{G} = \boldsymbol{E}_{\boldsymbol{MM}} + \boldsymbol{G}_{\boldsymbol{GB}} + \boldsymbol{G}_{\boldsymbol{SA}} - \boldsymbol{TS} \qquad \textbf{(2)}$$

$$\mathbf{E_{MM}} = \mathbf{E_{vdw}} + \mathbf{E_{ele}} + \mathbf{E_{int}} \qquad \textbf{(3)}$$

Where $E_{MM}$ is the molecular mechanical energy, $G_{GB}$ is polar contribution towards solvation energy calculated by Generalized Born (GB) method respectively. $G_{SA}$ is the contribution from nonpolar terms towards solvation energy, and TS is the entropic contribution of the inhibitor. $E_{MM}$ was obtained by summing contributions from, electrostatic energy ($E_{ele}$), vanderwaal energy ($E_{vdw}$), and internal energy including bond, angle, and torsional angle energy ($E_{int}$) using the same force field as that of MD simulations. $G_{GB}$ was calculated with Onufriev's method (Onufriev, Bashford & Case 2004)). $G_{SA}$ in equation 2 is proportional to the solvent accessible surface area (SASA) and was computed by molsurf module using the equation 5.

$$\boldsymbol{G}_{\boldsymbol{SA}} = \boldsymbol{\gamma} * \boldsymbol{SASA} + \boldsymbol{b} \qquad \textbf{(4)}$$

In equation 4, the surface tension proportionality constant (γ) was set to 0.0072 kcal/mol/$Å^2$, while the free energy of nonpolar solvation for a point solute (b) was set to a default value. SASA was computed by molsurf using a fast algorithm called the Linear Combinations of Pairwise Overlaps (LCPO), which simply computes a surface area of two overlapping atoms by subtracting their overlapping region from the total area of their vdW surfaces (Weiser, Shenkin & Still 1999)). vdW radii in AMBER and GAFF force fields were used to define solute atoms which were then augmented by a probe sphere of 1.4Å for the molsurf computations. Binding free energy calculations were averaged over 500 frames taken at the interval of 100 ps over the last 50 ns of 100 ns production run.

**Results and discussion**

To compare the binding free energies of NPV in PDE4B with, **U-Cat** complex, or without UCR2 domain *i.e.* **Cat complex**, we calculated MM-GB/SA energies of the two systems. The current comparative analysis involves a common inhibitor, NPV and we assumed that entropy have similar effects on both the similar complex structures. Therefore, we did not calculate the entropies for both the cases (**Table 1**).

The relative binding free energy ($\Delta G_{Total}$) of the **U-Cat** and **Cat** complex are -35.54 (± 2.23) kcal/mol and -24.57 (± 2.6) kcal/mol respectively. It is encouraging to find that these energies were consistent with our MM-GBSA studies.

The electrostatic ($\Delta \mathbf{E}_{Eel}$), van der Waals ($\Delta \mathbf{E}_{vdw}$), the nonpolar component of the solvation energy ($\Delta \mathbf{E}_{surf}$) and the polar component ($\Delta \mathbf{E}_{GB}$) of solvation free energy ($\Delta G_{solv}$), terms of the molecular mechanics energy were favourable for both the complex structures. The van der Waals component, $\mathbf{E}_{vdw}$ (**U-Cat**: -53.49 and **Cat**: -44.21 kcal/mol), and the nonpolar component of the solvation energy, $\mathbf{E}_{surf}$ (**U-Cat**: -6.65 and **Cat**: -5.52kcal/mol) showed the favorable contribution to the ligand protein complex formation. The total solvation energy, $\mathbf{\Delta G_{solv}}$ (**U-Cat**: -211.70 and **Cat**: -131.76 kcal/mol) is strongly favourable at both complexes. Although the hydrophobic contribution to solvation free energy is very less yet the polar component played very crucial role.

Electrostatic interactions were supposed to play a foremost major role in the ligand and a protein interaction. But it is of high importance to consider the electrostatic component of the molecular mechanics energy together with the polar contribution to solvation while deciding the role of electrostatics in the ligand-protein complex formation. At both protein-ligand complexes, the nature of the polar component of the total solvation energy is similar and thus favourable for the bound state of the ligand but the on the contrary the electrostatic contribution, $\mathbf{E}_{Eel}$ (**U-Cat**: 229.65 and **Cat**: 151.40 kcal/mol) does not play the expected significant role in the ligand complex interaction but this interaction were found to make a very small contribution to the binding free energy. The weakness of electrostatic interactions was surprising because of the conserved H-bond interaction between the invariant Gln615 and crystal ligands and reported inhibitors of PDE4B (Zhang et al. 2004;

Wang, Robinson & Ke 2007). Apart from the energy contribution to the complex-stability, like other analyses, here also we observed the effect of N-terminal. All the contributing energy and the binding free energy showed lower value for **U-Cat** in comparison to the **Cat** protein complex.

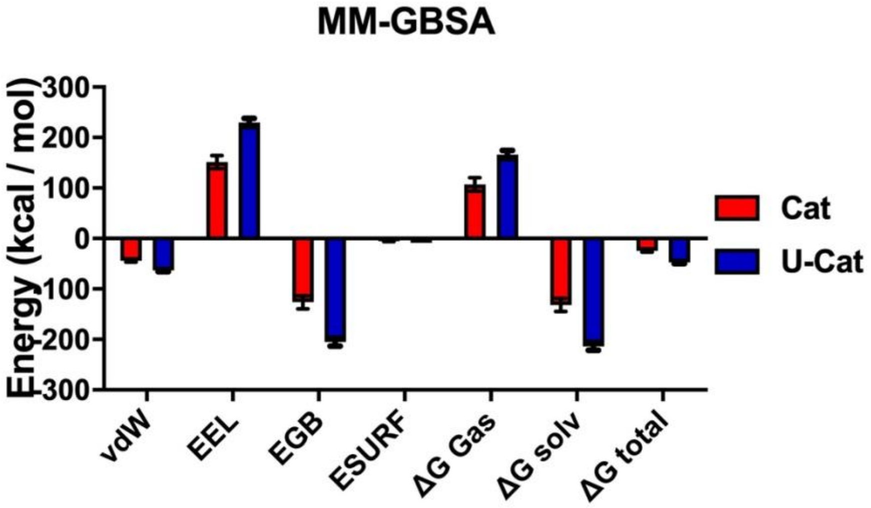


**Supplementray figure 1. MM-GBSA analysis of Cat and U-Cat versions of PDE 4B**
